# Metabolic network reductions

**DOI:** 10.1101/499251

**Authors:** Mojtaba Tefagh, Stephen P. Boyd

**Affiliations:** Information Systems Laboratory, Department of Electrical Engineering, Stanford University, Stanford, CA, USA

**Keywords:** systems biology, metabolic network analysis, canonical metabolic network reduction, quantitative flux coupling analysis, QFCA

## Abstract

Genome-scale metabolic networks are exceptionally huge and even efficient algorithms can take a while to run because of the sheer size of the problem instances. To address this problem, metabolic network reductions can substantially reduce the overwhelming size of the problem instances at hand. We begin by formulating some reasonable axioms defining what it means for a metabolic network reduction to be “canonical” which conceptually enforces reversibility without loss of any information on the feasible flux distributions. Then, we start to search for an efficient way to deduce some of the attributes of the original network from the reduced one in order to improve the performance. As the next step, we will demonstrate how to reduce a metabolic network repeatedly until no more reductions are possible. In the end, we sum up by pointing out some of the biological implications of this study apart from the computational aspects discussed earlier.

**Author summary:** Metabolic networks appear at first sight to be nothing more than an enormous body of reactions. The dynamics of each reaction obey the same fundamental laws and a metabolic network as a whole is the melange of its reactions. The oversight in this kind of reductionist thinking is that although the behavior of a metabolic network is determined by the states of its reactions in theory, nevertheless it cannot be inferred directly from them in practice. Apart from the infeasibility of this viewpoint, metabolic pathways are what explain the biological functions of the organism and thus also what we are frequently concerned about at the system level.

Canonical metabolic network reductions decrease the number of reactions substantially despite leaving the metabolic pathways intact. In other words, the reduced metabolic networks are smaller in size while retaining the same metabolic pathways. The possibility of such operations is rooted in the fact that the total degrees of freedom of a metabolic network in the steady-state conditions are significantly lower than the number of its reactions because of some emergent redundancies. Strangely enough, these redundancies turn out to be very well-studied in the literature.

## 1 Introduction

Two decades ago, the first genome-scale metabolic network reconstruction of the cellular metabolism of an organism was published [1] shortly after the first genome was sequenced. From that time on, the ever-increasing advances in the high-throughput omics technologies have allowed for the more and more comprehensive reconstructions of exponentially growing sizes [2, 3, 4, 5, 6, 7, 8, 9, 10, 11, 12, 13, 14, 15, 16, 17, 18, 19]. These reconstructions have numerous applications in contextualization of high-throughput data, guidance of metabolic engineering, directing hypothesis-driven discovery, interrogation of multi-species relationships, and network property discovery [20]. However, the vast amount of data for some organisms can be a two-edged sword which makes many essential tasks in the metabolic network analysis computationally intractable.

To overcome the demands of systems biology, even while they are outpacing Moore’s law [21], faster computational techniques are needed to enable the current methods to scale up to match the progress of high-throughput data generation in a prospective manner. As a natural solution, reducing the size of genome-scale metabolic networks has always been exploited to the advantage of performance for predetermined computational tasks [22, 23, 24, 25, 26]. However, all of these mentioned studies assume that a set of protected reactions which must be retained in the reduced metabolic network is given in advance and the other reactions are dispensable. In this way, these metabolic network reductions are not agnostic to the downstream analysis and lose information on some of the reactions which are irrelevant to the task at hand.

In the field of *flux coupling analysis* (FCA), from the very beginning *Flux Coupling Finder* (FCF) [27] considers aggregating all the isozymes and removing the blocked reactions. More recently, *fast flux coupling calculator* (F2C2) [28] considers merging the enzyme subsets [29] too. *FluxAnalyzer* [30] also detects conservation relations as a preprocessing step. *MONGOOSE* [31] employs a similar loss-free network reduction to convert the input model into a canonical form.

Although the practical implications of these findings affirm the plausibility of their general-purpose reductions in a common sense approach, to the best of our knowledge the formal definition of a metabolic network reduction, in general, is not clear so far. The aforementioned studies provide strong support for the idea that such general-purpose reductions preserve some interesting attributes of metabolic networks, *e.g.*, the flux coupling features. In this article, we specify a broad class of metabolic network reductions with respect to which some generally desired properties of metabolic networks are invariant.

### Outline

In §2, we briefly overview the FCA framework. In §3, we define and study single and multiple sequential metabolic network reductions, respectively. Afterward in §4, we argue that the reduced metabolic networks are not only interesting from the computational perspective, but also they are biologically interpretable. Finally in §5, we conclude by highlighting the major contributions of this work.

## 2 Background

We specify a metabolic network by an ordered quadruple 𝒩 =(ℳ, ℛ, 𝒮, ℐ).where 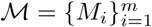 denotes the set of metabolites of size *m*,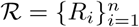 denotes the set of reactions of size *n*, 𝒮 denotes the *m × n* stoichiometric matrix, and ℐ*⊆* ℛ denotes the set of irreversible reactions. Since it is often the case that *n » m*, we consider *n* as the size of 𝒩 too.

In the constraint-based analysis of metabolic networks, the constraints 𝒮 *v* = 0 and *v*_*I*_*≥*0 are imposed on the metabolic network by the steady-state conditions and the definition of irreversible reactions, respectively. By a slight abuse of notation, *v*_*I*_*≥*0 means *v*_*i*_*≥*0 for all the indices *i* for which *R*_*i*_ *∈*ℐ We denote the *steady-state flux cone* [32] *by*

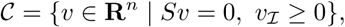

The feasible flux distributions of 𝒩 are then defined to be the members of 𝒞.

We call *R*_*k*_ *∈*ℛ a blocked reaction if *v*_*k*_ = 0 for all the feasible flux distributions *v* ∈ 𝒞. To the end of this paper, whenever we assume that all the blocked reactions are removed, by this, we also assume that if a reversible reaction *R*_*k*_ is blocked in only one direction, then this blocked direction is removed too meaning that *R*_*k*_ is included in ℐ subsequently. In §3, we will review these trivial ways of reducing a metabolic network in more details.

In order to derive even more metabolic network reductions, we should exploit other redundancies of the steady-state flux cone analogous to the case of blocked reactions where the rate of a blocked reaction is always zero irrespective of what the other flux coefficients are equal to. A somewhat less obvious situation is when the rate of one reaction is unambiguously determined by another one as is the case in the following definitions of FCA.

**Definition 1** ([27]). For an arbitrary pair of unblocked reactions *R*_*i*_, *R*_*j*_ *∈*ℛ:

### Directional coupling

*R*_*i*_ is *directionally coupled* to *R*_*j*_, denoted by *R*_*i*_ *→ R*_*j*_, if for all feasible flux distributions *v*_*i*_ ≠ 0 implies *v*_*j*_ ≠ 0.

### Partial coupling

*R*_*i*_ is *partially coupled* to *R*_*j*_, denoted by *R*_*i*_ *←→ R*_*j*_, if both *R*_*i*_ *→ R*_*j*_ and vice versa *R*_*j*_ *→ R*_*i*_ hold.

### Full coupling

*R*_*i*_ is *fully coupled* to *R*_*j*_, denoted by *R*_*i*_⟺*R*_*j*_, if there exists a *full coupling equation* (FCE)

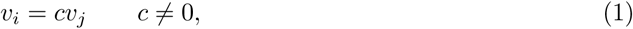

which holds for all *v ∈𝒞*.

In this definition, full coupling is defined by means of a flux coupling equation, but directional and partial couplings are defined qualitatively. We can apply the following proposition to get the equivalent quantitative definitions of directional and partial couplings by introducing two other flux coupling equations in analogy to FCE. *Quantitative flux coupling analysis* (QFCA) [33] redefines the original flux coupling relations of FCA by the existence of flux coupling equations.

**Proposition 1** ([33]). *Suppose that* 𝒩= (ℳ,ℛ,𝒮,ℐ) *has no irreversible blocked reactions. Let R*_*j*_ *be an arbitrary unblocked reaction, and* 𝒟_*j*_ *⊆*ℐ *denote the set of all the irreversible reactions which are directionally coupled to R*_*j*_ *excluding itself. Then*, 𝒟_*j*_ *≠ ∅ if and only if there exists c*_*d*_ *>* 0 *for each R*_*d*_ *∈*𝒟_*j*_, *such that the following* directional coupling equation *(DCE)*

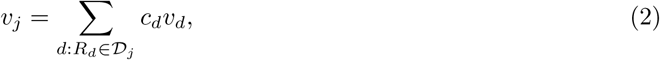

*holds for all v* ∈ 𝒞. *Moreover, for any unblocked R*_*i*_ ∉, ℐ *we have R*_*i*_ *→ R*_*j*_ *if and only if there exists an* extended directional coupling equation *(EDCE)*

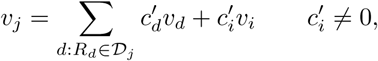

*which holds for all v ∈*𝒞.

Prior to QFCA, the general attitude towards directional coupling was that it is a one-way relation which is, of course, true considering only one pair of reactions. However QFCA, for the first time, revealed that this relation becomes two-sided if we consider the set of directionally coupled reactions to a single reaction which are sufficient to infer the flux coefficient of it. This redundancy motivates generalizing the previously known application of FCE in the metabolic network reductions to DCE.

In the sequel, we will formally show that metabolic network reductions are intrinsically related to QFCA. The intuition is that redundancy in the steady-state flux cone creates flux coupling relations, and metabolic network reductions reduce redundancy by compressing this information. Identifying and eliminating statistical redundancy is called lossless compression in information theory [34]. We will prove results of the same flavor to show that QFCA induces lossless reductions and any other metabolic network reduction beyond that is a lossy compression.

We continue by the following general mathematical definition, and subsequently we use this terminology to formalize the notion of a minimal feasible flux distribution. The support of a flux distribution *v∈* **R**^*n*^ is denoted by supp(*v*), and is defined to be the set of those reactions in **ℛ** which are active in this flux distribution, *i.e.*,

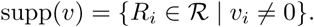

**Definition 2** ([35]). We call a nonzero feasible flux distribution 0 ≠ *v* ∈ 𝒞 an *elementary mode* (EM), if its support is minimal, or equivalently, if there does not exist any other nonzero feasible flux distribution 0 ≠ *u∈*𝒞 such that supp(*u*) *⊂* supp(*v*).

Given a finite number of vectors *v*_1_, *v*_2_, *…, v*_*d*_, a *conic combination* of these vectors is any weighted sum *θ*_1_*v*_1_ + *θ*_2_*v*_2_ + *…* + *θ*_*d*_*v*_*d*_ with nonnegative weights *θ*_1_, *θ*_2_, *…, θ*_*d*_ *≥* 0. The set of all such conic combinations is called the polyhedral convex cone generated by *v*_1_, *v*_2_, *…, v*_*d*_ [36].

One of the main properties of EM is that the set of all EMs in a metabolic network generates its steady-state flux cone. Another way of expressing this is to say that any arbitrary feasible flux distribution can be written as a conic combination of EMs. With QFCA in mind, this property implies that to establish a flux coupling relation instead of going through all the feasible flux distributions, it suffices to validate the corresponding flux coupling equation only for the EMs. Accordingly, reducing the size of a metabolic network so long as preserving the information on its EMs and in turn, the flux coupling relations leads to a significant speed-up during QFCA.

As we will see in the next section, the relationship between metabolic network reductions and QFCA does not end here and is surprisingly reciprocal. The uncovered connection between these two concepts sheds light on the mutual nature of both in relation to the core concept of EM. Besides presenting an axiomatic characterization to rigorously examine this connection, we will also derive metabolic network reductions inspired by QFCA which, though intuitive and straightforward, are nevertheless general enough to reduce any arbitrary metabolic network into a minimal irreducible one.

## 3 Methods

As an illustrative example of the method, consider the following toy metabolic network in Fig 1. We name it 𝒩 = (ℳ, ℛ, 𝒮, ℐ), where ℳ = {*M*_1_, *M*_2_, *M*_3_}, ℛ = {*R*_1_, *R*_2_, *R*_3_, *R*_4_, *R*_5_}, ℐ = ℛ, and

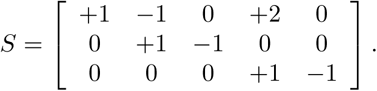

Note that, *R*_2_ ⟺ *R*_3_ by the corresponding FCE *v*_2_ = *v*_3_ for all *v* ∈ 𝒞, which we will represent by 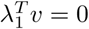 where 

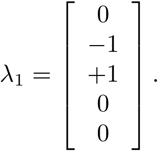

Therefore, for all *v* ∈ 𝒞

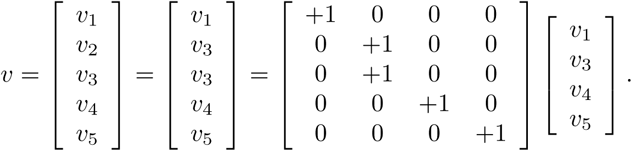

Let *ϕ*_1_ denote the matrix on the right hand side. For all *v* ∈ **R**^5^,

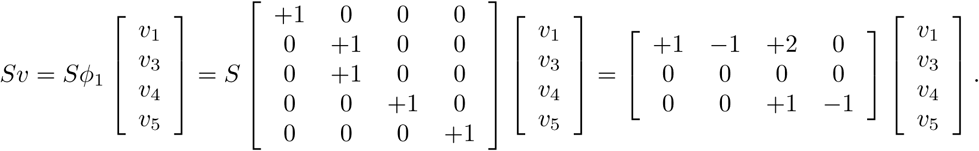

As a result, if we define

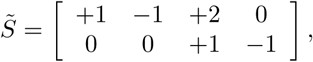

then we have that *v* ∈ 𝒞 if and only if

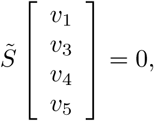

and *v*_*i*_ *≥* 0 for *i* = 1, 3, 4, 5.

**Figure 1:**
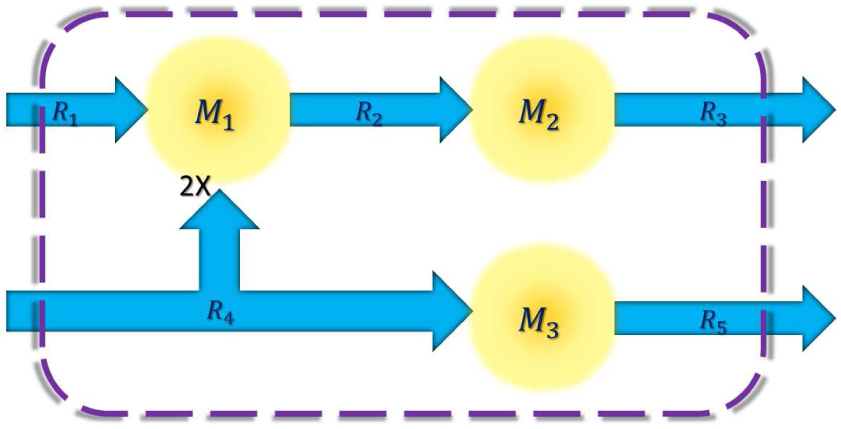
The original metabolic network

Intuitively, if we incorporate *R*_2_ into *R*_3_ and remove *M*_2_, then we arrive at the reduced metabolic network in Fig 2, namely 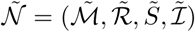, where 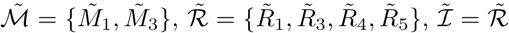,and 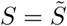.

**Figure 2:**
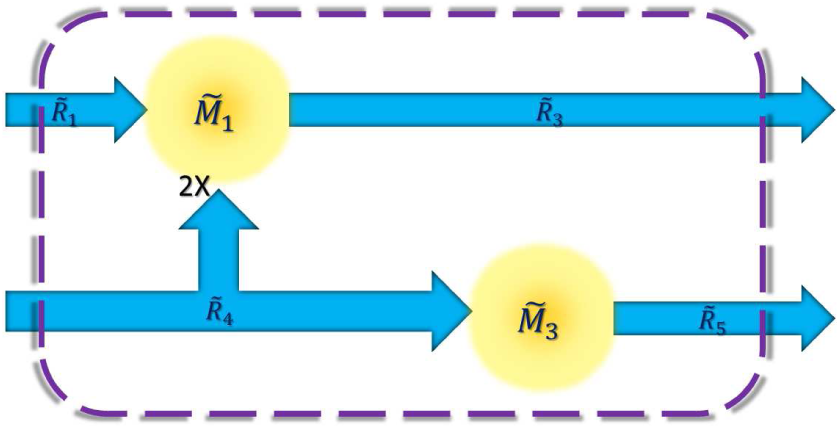
The reduced metabolic network

Similar to the previous case, *R*_1_, *R*_4_ → *R*_2_ by the corresponding DCE *v*_2_ = *v*_1_ + 2*v*_4_ for all *v* ∈ 𝒞, which we will represent by 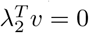 where

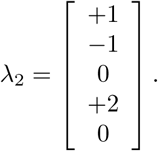

Similarly, we define *Ø*_2_ such that

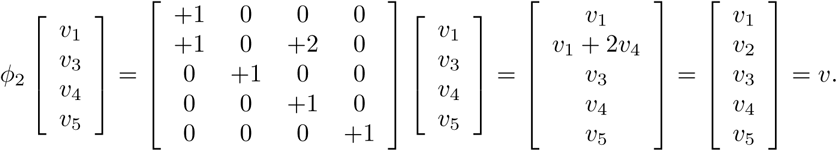

Therefore,

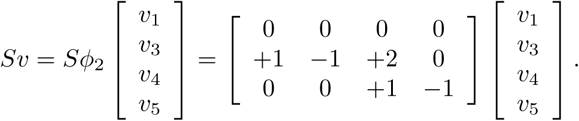

Comparing 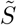 with *S ϕ*_2_ they are equal if we remove the row corresponding to *M*_1_. We will come back to this point for a more careful inspection in *§*4, however for now, removing an all zeros row from the stoichiometric matrix does not affect the steady-state flux cone, hence, we can safely do it.

This time, if we incorporate *R*_2_ into *R*_1_ and *R*_4_ and remove *M*_1_, then we arrive at the reduced metabolic network in Fig 3, which is the same 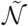 except that 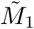 is replaced by 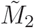 which again, does not affect the steady-state flux cone.

**Figure 3:**
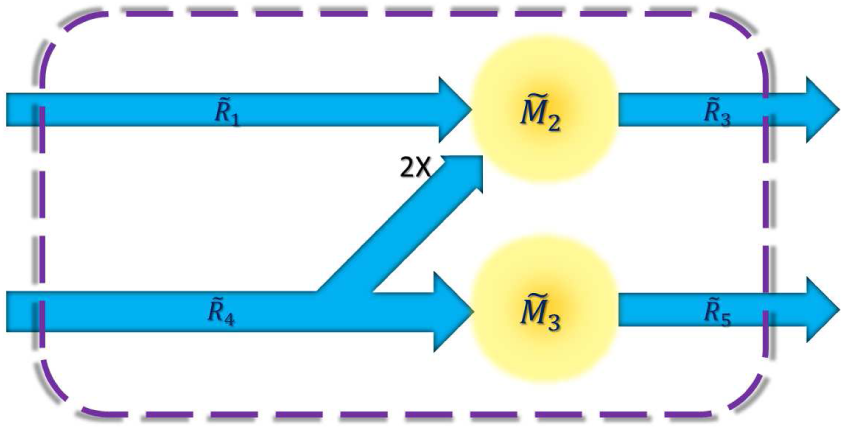
A DCE-induced reduction

Having said that, recall the correspondence between the original and reduced reactions in both cases. For the FCE-induced reduction we had,

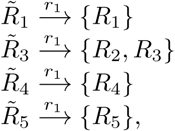

but for the DCE-induced reduction we had,

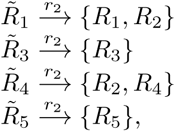

where we call *r*_*i*_ the reduction map. Note that *r*_*i*_ is uniquely determined by the flux coupling relations in both cases.

Let *e*_*k*_ denote the *k*th unit vector of the standard basis of **R**^5^. Comparing *ϕ*_*i*_ with the following matrix

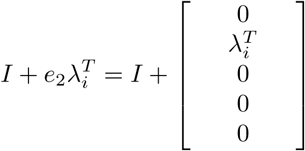

they are equal for both *i* = 1, 2 if we remove the all zeros column corresponding to *R*_2_. Hence, *ϕ*_*i*_ is uniquely determined by the flux coupling equations in both cases.

From now on, suppose λ is an arbitrary flux coupling equation for *R*_2_, *i.e.*,

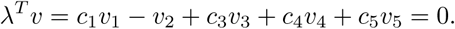

For an index *i*, let *X*^*i*^ and *x*^*i*^ denote the *i*th column of the matrix *X* and the *i*th entry of the vector *x*. Furthermore, let *X*^(*i*)^ and *x*^(*i*)^ denote the result of removing *X*^*i*^ and *x*^*i*^ from *X* and *x*, respectively. By this notation, we define

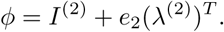

Like before, *ϕ* maps the reduced *v*^(2)^ to the original *v*.

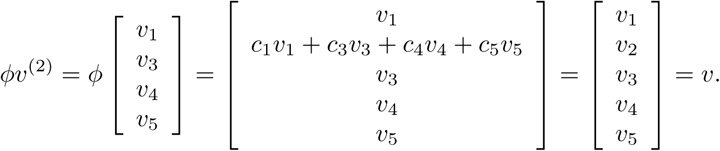

Moreover,

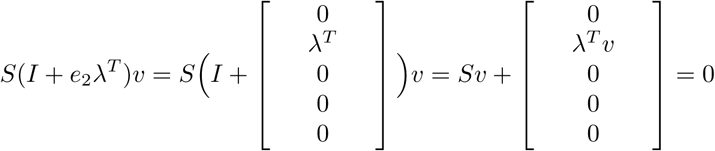

for all *v* ∈ 𝒞. However, the second column of *I* + *e*_2_λ^*T*^ is zero so we can remove it together with the second entry of *v* to arrive at the equation *Sϕv*^(2)^ = 0. Hence, *ϕ* also provides the reduced stoichiometric matrix by setting 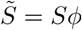.

Furthermore, the reversibility of the reduced reactions is determined by the simple rule that 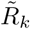 is irreversible if and only if 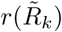 contains at least one irreversible reaction, and the reduction map *r* is uniquely determined by which flux coupling coefficients are nonzero in λ.

The last three paragraphs can be generalized from this toy metabolic network with five reactions to the most general case by following the same steps. In summary, we remove the blocked reactions, then merge the coupled reactions by adding the corresponding columns of the stoichiometric matrix and making the merged reaction irreversible if any of the original ones were irreversible. In the Appendix, we prove that all these QFCA reductions preserve the set of EMs. Moreover, we have abstracted the properties of these reductions used in the proof to show that this result holds for not only the QFCA reductions, but also for a broad class of reductions which we call canonical metabolic network reductions.

EM analysis, while still limited by the size of the metabolic networks it can handle, is a powerful metabolic pathway analysis tool for a wide variety of application such as characterizing cellular metabolism and reprogramming microbial metabolic pathways [37, 38, 39]. This potential encourages conceptualizing the notion of a canonical metabolic network reduction which respects EMs as has been proposed by [30] long ago. In order to realize this idea, we will propose three axioms for characterizing a canonical metabolic network reduction.

Let 𝒩 = (ℳ, ℛ, 𝒮, ℐ) be an arbitrary metabolic network. We say that the metabolic network 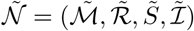 is a reduction of 𝒩 if

1. there exists a surjection *ϕ*: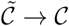, where 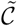 and 𝒞 are the corresponding steady-state flux cones;
2. there exists a reduction map 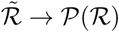 such that for any 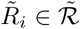 we have

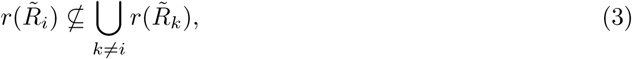

where 𝒫 (ℛ) denotes the power set (the set of all the subsets) of ℛ;
3. and the following diagram commutes

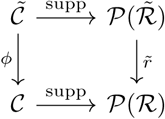

where 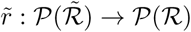 is defined by

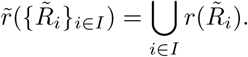

We have dedicated an entire Appendix to investigate the properties of such canonical metabolic network reduction. However, to continue reading the rest of the paper, we only need the following corollary of the reduction theorem which is proved in the Appendix.

**Corollary 3.0.1.** *By setting i* = *j in the reduction theorem, any reaction in* 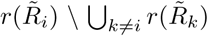 *is directionally coupled to any reaction in* 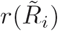.

The first axiom enforces that each feasible flux distribution in either the reduced or the original metabolic network corresponds to at least one feasible flux distribution in the other one. The second axiom introduces the mapping *r* between reactions, which induces the injection 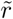 between pathways essential for the third axiom to enforce that having minimal support is an invariant property under *ϕ*. This, in turn, implies that the EMs of these two metabolic networks are in correspondence with each other.

The only suspect property in here which seems like a strong assertion is (3) in the second axiom. Recall that by Corollary 3.0.1, all the reactions in 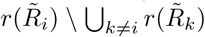 turn out to be fully or partially coupled to each other, and hence, are intimately related. Consequently, instead of some abstract 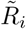 in an unreal metabolic network, we can think of each 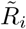 as a class of fully or partially coupled reactions in the current metabolic network which is very similar to the notion of enzyme subsets [29]. Whereas, if for some 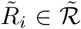 we had 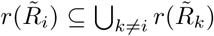, then no reaction in ℛ was uniquely associated to 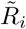. To sum up, the main reason behind this assertion is to ensure that reactions in 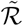 are meaningful from the biological point of view.

We give the example of coupled metabolites to further demonstrate this point. *Metabolite concentration coupling analysis* (MCCA) [40] is the same as FCA except it is applied to the left null space of the stoichiometric matrix [41]. We refer the interested reader to the results of [40] for an extensive discussion of the biological relevance of the coupled metabolites. Note that the subsets 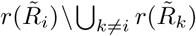 are exactly the same metabolite subsets in this instance. Furthermore, the reduced metabolic network preserves the reactions among these metabolically meaningful pools in addition to all the MCCA results derived from FCA. These reactions restrain the minimal conserved pools (the analogue of elementary modes) to be in one-to-one correspondence with the original set for *minimal conserved pool identification* (MCPI) [40].

Let 𝒩 = (ℳ, ℛ, 𝒮, ℐ) be an arbitrary metabolic network, and 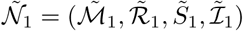 be a reduction of 𝒩 by *ϕ*_1_ and *r*_1_. Suppose that we apply several metabolic network reductions to 𝒩 successively. For instance, suppose that we reduce 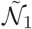 once more to get 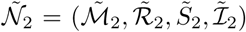 by a different metabolic network reduction, namely the one corresponding to *ϕ*_2_ and *r*_2_. We will show that the composition of consecutive reductions is again a reduction itself.

1. *ϕ*_1_ ° *ϕ*_2_ : 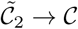 is a surjection because the composition of surjective functions is surjective;
2. 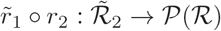 is a legitimate reduction map because for any 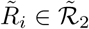 we have

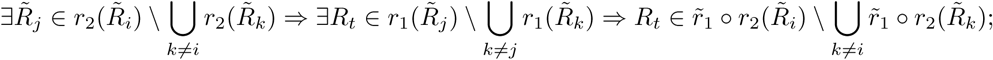
3. and the following diagram commutes

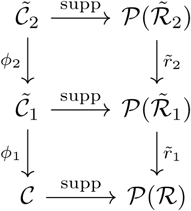

because for any 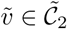

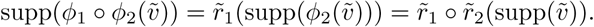

Putting it all together, 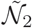 is a reduction of 𝒩 by the surjection *ϕ*_1_ ∘ *ϕ*_2_ and the reduction map 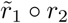.

Up to now, we have covered how to reduce the metabolic network by the following three stages.

- First, we eliminate all the blocked reactions.
- Second, we merge all the fully coupled reactions.
- Third, we remove the eligible reactions by the DCE-induced reductions.

Suppose that after conducting all these three stages, the last metabolic network is 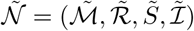 of size *ñ*, thus we obtain the chain of reductions

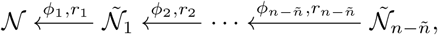

where 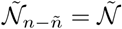, since each reduction reduces exactly one reaction at a time. Throughout this process, we consider the aggregated metabolic network reduction resulting from combining the previously applied ones up to some point in time as a single reduction, *i.e.*,

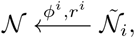

where *ϕ*^*i*^ = *ϕ*_1_ ∘ … ∘ *ϕ*_*i*_ and 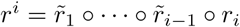.

We claim that each *ϕ*^*i*^ is a one-to-one correspondence when restricted to the EMs. In the Appendix, we prove that 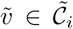 is an EM if and only if 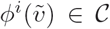 is an EM. Furthermore, *ϕ*^*i*^ is surjective by definition. It only remains to show that *ϕ*^*i*^ is injective too. One can easily verify the injectivity of the single metabolic network reductions *ϕ*_*t*_ of any of these three kinds separately. Hence, it is also proved for *ϕ*^*i*^ which is the composition of *i* of them.

As a consequence, the overall reduction generated by compositing all the preceding three stages is a metabolic network reduction which provides a one-to-one correspondence between the EMs of 𝒩 and 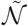. This can also be shown independently, given the straightforward form of this specific reduction which we characterize next.

For the sake of simplicity, assume that we arrange ℛ in an ordering such that first come the blocked reactions removed in the first stage, ℬ = {*R*_1_, *…, R*_*b*_}. After ℬ come the reactions which persist to the end, and by this we mean the final set 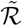 which is a subset of ℛ, because 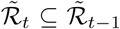 for each of these three types of reductions individually. Moreover, assume that 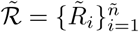 is sorted in such a way that 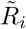 is fully coupled to *a*_*i*_ other reactions in ℛ where *a*_1_ ≥ *a*_2_ ≥ … ≥ *a*_*f*_ *> a*_*f*+1_ = = *a*_*ñ*_ = 0 (the descending order of the number of fully coupled reactions in). At last come the reactions which were removed in stages two and three, respectively.

By these simplifying assumptions, if *P* is the permutation matrix which permutes the columns of *S* to the ordering we have assumed, then we can give the following explicit formulas for 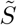, *ϕ*^*n-ñ*^, and *r*^*n-ñ*^ which is 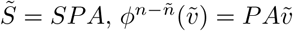, and 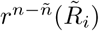 is equal to the index set of the nonzero entries of the *i*th column of *A* with the following structure (see Fig 4).

**Figure 4:**
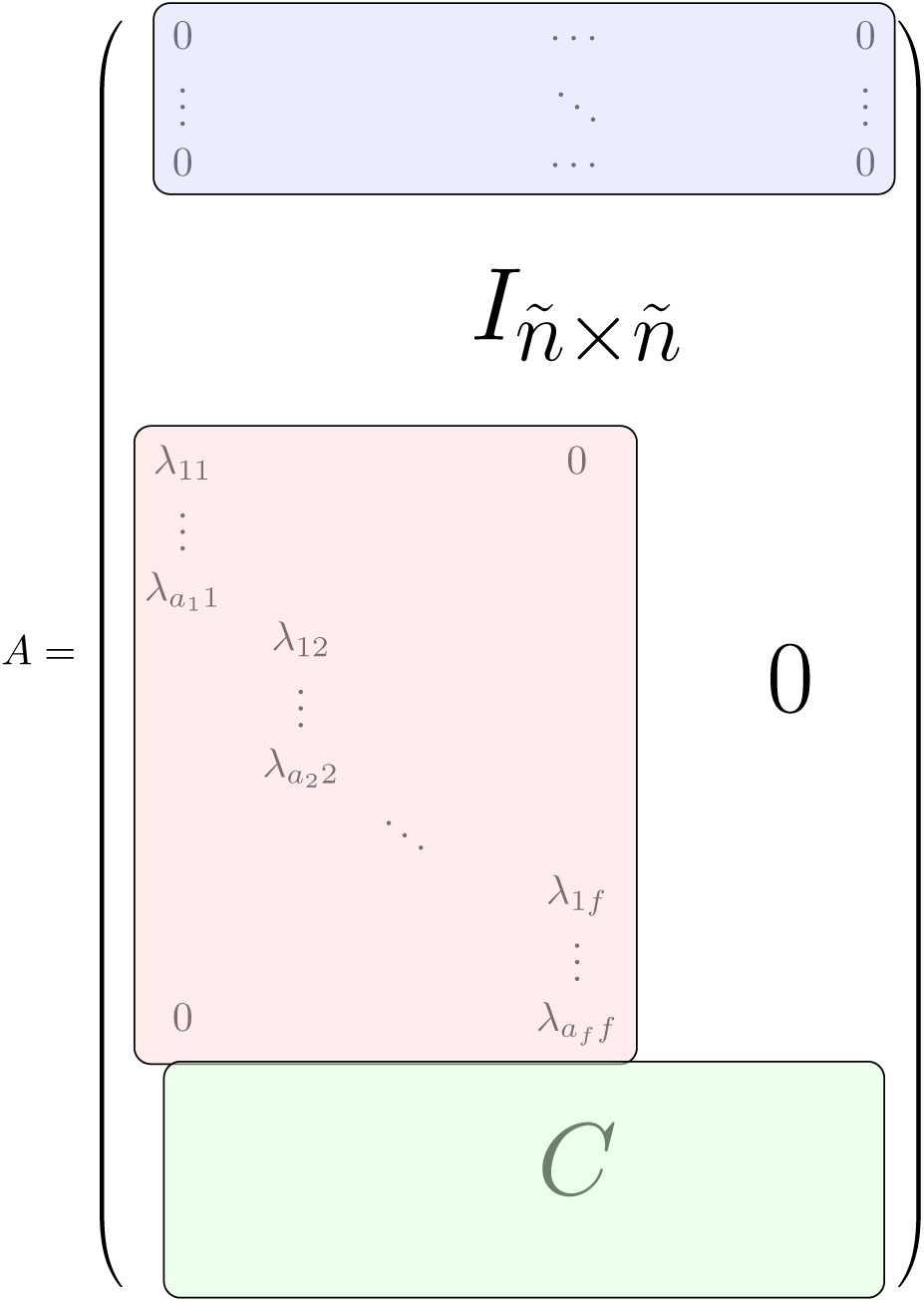
Sparsity pattern of A

As depicted in Fig 4, the blue block has *b* zero rows corresponding to the blocked reactions which always have zero flux coefficients. The identity matrix corresponds to 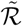 which in turn is associated with a subset of ℛ. The red block corresponds to the fully coupled reactions removed in stage two, and its nonzero entries are the associated FCE coefficients, *i.e., c* in (1). In the end, each row of the nonnegative matrix *C* corresponds to a reaction removed in stage three, and its positive entries are the associated DCE coefficients, *i.e., c*_*d*_ in (2).

Ultimately by Corollary 3.0.1, because of the fact that 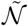 has no directionally coupled pair of reactions, otherwise we could have continued stage three, the size of 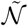 cannot be reduced any further. Even more than that, not only 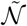 is minimal in the sense that it cannot be reduced any further without violating one of the principles we have assumed for the canonical metabolic network reductions, but also it has the minimum size, and in this sense, this is the most compact form of an equivalent steady-state flux cone with identical EMs.

For the proof of this statement, we show that no two distinct reactions in 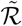 can be reduced to the same reaction in any possible metabolic network reduction, hence *ñ* is the minimum size. Let 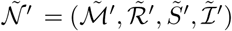 be an arbitrary reduction of 𝒩 by *ϕ*′ and *r*′. By Corollary 3.0.1, if 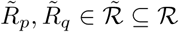 are reduced to the same reaction 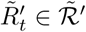, then there exists 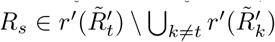 which is directionally coupled to both 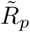 and 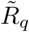, simultaneously. However, 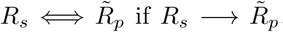, otherwise 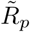 would have been removed in the third stage. Therefore 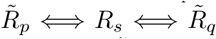, which cannot hold unless *p* = *q* since stage two guarantees that no two distinct reactions in 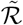 are fully coupled to each other.

## 4 Discussion and results

QFCA^1^ is implemented in the COBRA toolbox v3.0 [42], publicly available for academic and research purposes. This MATLAB^©^ package is also the first implementation of the reduction algorithm presented here. It paves the way for the development of scalable solutions for various computational tasks. This brings us to a suggestion for the future research to work out the details of integrating this universal preprocessing (or postprocessing as in *NetworkReducer* [25]) step with the compatible pipelines of interest.

The reduced stoichiometric matrix introduced in §3 has a great potential to speed-up the down-stream analyses which only rely on EMs, in particular, QFCA. The smaller the reduced 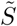, the faster any downstream analysis on this proxy stoichiometric matrix. For a running example, iMM1415 (*Mus Musculus*, 1415 genes) [43] has 3726 reactions initially, and MONGOOSE reduces it into 1625 reactions while QFCA achieves a record 1171 reactions in the same way as we described in §3.

Even though 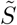 has *m* rows in theory, it turns out that for almost every real-world metabolic network, many of its rows are identically zero and therefore, for all practical purposes they may be removed safely. For instance, consider the metabolic network in Fig 1 and suppose that we remove *R*_2_ by the corresponding DCE-induced reduction. Although, *M*_1_ is still there after conducting the metabolic network reduction according to the procedure of §3, it can be removed to arrive at the metabolic network in Fig 3 since it is no longer involved in any reactions.

Moreover, we remove the linearly dependent rows of 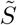 as described under the title of the detection of conservation relations by [30]. In practice,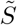 may be much smaller than *S* both regarding the number of rows and columns. Going back to our running example, the number of metabolites can be reduced from 2775 to 335 by QFCA while the corresponding number for MONGOOSE is 643.

We should mention that in spite of the fact that the reduced 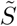 has a smaller size than the original *S*, it might be denser for some instances. As a result, if some algorithm is already exploiting the sparsity of *S* intricately, perhaps it does not gain much from this preprocessing step. This is a common theme in numerical analysis that sometimes sparse computation can make up for not including off-the-shelf preprocessing steps reducing the size of sparse matrices. Nonetheless, in our running example, the number of nonzero elements of *S* and 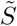 are 14053 and 14385, respectively. These numbers are promising as they indicate that the reduced stoichiometric matrix is nearly as sparse as the original one.

To showcase canonical metabolic network reductions on a large-scale reference database, we have reduced the BiGG universal model [44] of 13249 metabolites, 24311 reactions, and 95774 nonzero stoichiometric coefficients into a metabolic network with 1278 metabolites, 10255 reactions, and 56457 nonzero stoichiometric coefficients. Furthermore, we have reconstructed consistent metabolic networks for randomly sampled core sets of varying sizes, from zero up to the number of unblocked reactions, by applying the algorithm swiftcore [45] to both of these models. The runtime on the reduced network is 29% of the runtime on the original network averaged over 100 iterations (see Fig 5). Both results for each set of core reactions are double checked and are always flux consistent in both cases.

**Figure 5:**
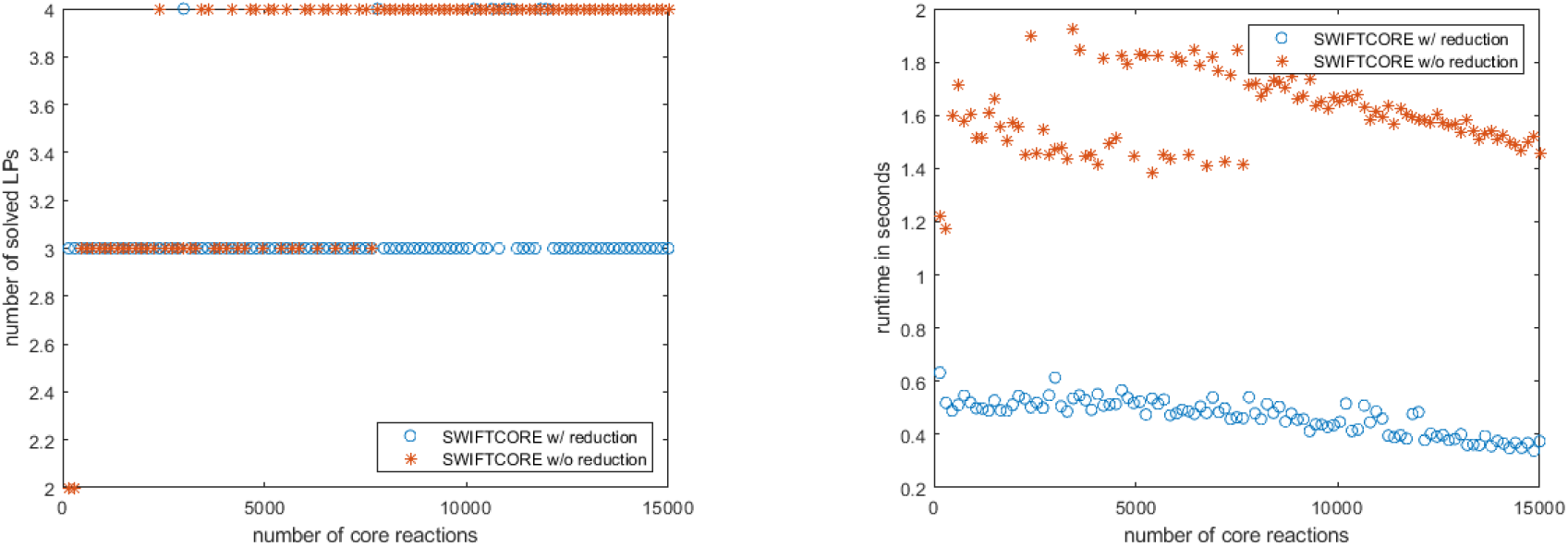
SWIFTCORE runs more than 3× faster on the reduced BiGG universal model

Apart from the computational benefits of canonical metabolic network reductions, the fact that the redundancies removed by the axiomatic framework coincide with the well-known biological QFCA relations is worth special attention. Recall that the reactions reduced during the suggested process are the reactions which the rest of the remaining reactions are coupled to. A recent study [46] has shown that these are the essential reactions enriching the vital metabolic processes of the cell. In addition, they are more conserved and of older evolutionary age. These reactions are essential in a wide range of conditions, and their associated genes are more expressed [47]. All these observations are compatible with the fact that canonical metabolic network reductions only keep the reactions which discriminate different elementary modes by their distinct activity patterns. For example, the first exchange reactions in pathways are merged into the intermediate reactions which differentiate those pathways from each other which is in agreement with the findings of [46].

## 5 Conclusions

Metabolic networks contain many essential reactions which are active regardless of the others. On the other hand, there are reactions which are specific to a few elementary modes and whose activity patterns determine the active pathway. Metabolic networks can be reduced by integrating the former reactions into the latter ones without mixing the elementary mode.

We have derived the minimum reduced metabolic networks by QFCA. Afterward, we have also proved a converse that any further reduction loses some information about the support of active pathways in analogy to the concept of lossy compression in information theory.

### Appendix

Let *ϕ*^-1^(*v*) denote the preimage of *v* ∈𝒞 under *ϕ*. The term supp(*ϕ*^-1^(*v*)) is well-defined as for any 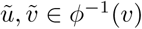, we have

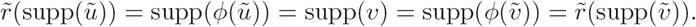

Therefore, 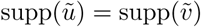 since 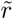 is injective from (3).

Another observation is that for any 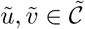

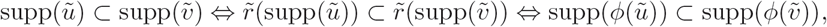

which is deduced from the second and third properties of the metabolic network reductions, respectively.

We claim that if 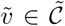 is an EM, then 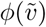 is also an EM. The proof is by contradiction. Let us suppose that there exists *u* ∈ 𝒞 such that:

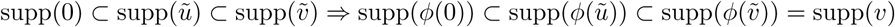

Since 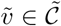 is an EM, for any 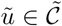 if 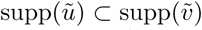, then supp(ũ) = ∅. However, supp(*ϕ*^-1^(0)) = supp(*ϕ*^-1^(*u*)) = ∅ is in contradiction with supp(*ϕ*^-1^(0)) ⊂ supp(*ϕ*^-1^(*u*)) which proves the desired result.

For the converse, we claim that if *v* ∈ 𝒞 is an EM, then any arbitrary 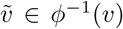 is an EM too. Assume to the contrary that for a fixed 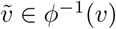 there exists 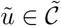 such that:

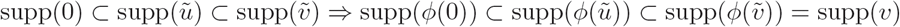

Since *v* ∈ 𝒞 is an EM, for any *u* ∈ 𝒞if supp(*u*) ⊂ supp(*v*), then *u* = 0. However, *ϕ*(0) = *ϕ*(ũ) = 0 is in contradiction with supp(*ϕ*(0)) ⊂ supp(*ϕ*(ũ)) which completes the proof.

Altogether, for any arbitrary metabolic network reduction 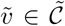 is an EM if and only if 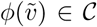 is an EM. As mentioned earlier, it is possible to infer flux coupling relations from EMs, solely. This evidence suggests that one should be able to deduce some of the flux coupling relations of the original and reduced networks from one another. The following theorem demonstrates one of the many possible ways to do so.

**Theorem 5.1** (The reduction theorem). *Suppose that* 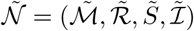 *is a metabolic network reduction of* 𝒩 = (ℳ, ℛ, 𝒮, ℐ) *by the surjection ϕ* : 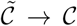 *and the reduction map r* : 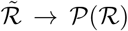. *For each* 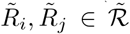 *such that* 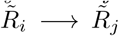, *any reaction in* 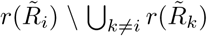 *is directionally coupled to any reaction in* 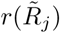, *Conversely, if there exists a reaction in* 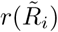 *which is directionally coupled to some reaction in* 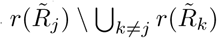. *then* 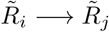.

*Proof.* In order to show the forward direction of the theorem, assume that 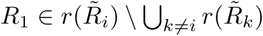 and 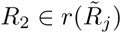. If *v*_1_ ≠ 0 for some arbitrary *v* ∈ *C*, then we have

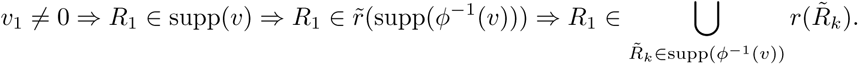

From the assumption that 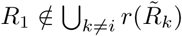, we have

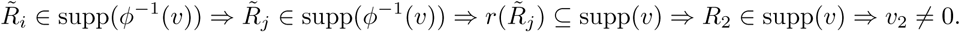

Since *v* ∈ 𝒞 was arbitrary, this proves one direction of the theorem.

To show the other direction of the theorem, assume that 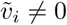 for some arbitrary 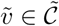. Additionally, assume 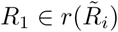 and 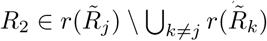 are such that *R*_1_ → *R*_2_. Then,

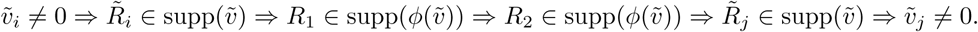

Therefore,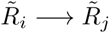 as was desired. □

As an introductory example, suppose that in 𝒩 = (ℳ, ℛ, 𝒮, ℐ), *R*_*i*_ is blocked. Let *S*^*i*^ and *S*^(*i*)^ denote the *i*th column of *S* and the resulting submatrix after removing it from *S*, respectively. Trivially, 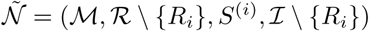 is a reduction of 𝒩 by letting

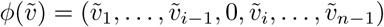

and *r*(*R*_*d*_) = {*R*_*d*_} for all *d* ≠ *i*. In a similar manner, if *R*_*i*_ ∉ ℐ and it is only blocked in the reverse direction, then 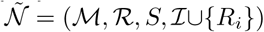 is a reduction of 𝒩 by letting 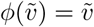 and *r*(*R*_*d*_) = {*R*_*d*_} for all *d*. On the other hand, if *R*_*i*_ ∉ ℐ and it is only blocked in the forward direction, then 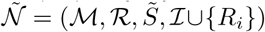 where

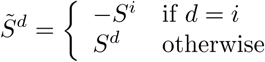

is a reduction of 𝒩 by letting

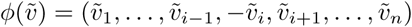

and *r*(*R*_*d*_) = {*R*_*d*_ }for all *d*. By applying the reduction theorem, 𝒩 and 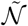 have exactly the same set of flux coupling relations in all these three cases.

For another example, suppose that in 𝒩 = (ℳ, ℛ, 𝒮, ℐ), distinct *R*_*i*_ and *R*_*j*_ are fully coupled with (1) as their corresponding FCE. If we define 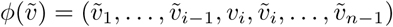 where

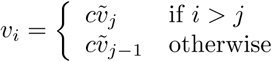

for the surjection, and

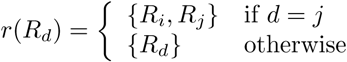

for the reduction map, then the metabolic network specified by 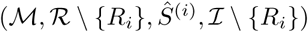, where

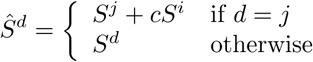

is a reduction of 𝒩.

By applying the reduction theorem, all of the flux coupling relations of this reduced network are valid in the original network as well. Furthermore, if in the reduced network *R*_*k*_ is directionally, partially, or fully coupled to *R*_*j*_ or vice versa, then in the original network *R*_*k*_ is directionally, partially, or fully coupled to *R*_*i*_ or vice versa, and these are all the additional flux coupling relations of 𝒩.

Arguably, fully coupled reactions are the most comprehensible ones as FCE gives an entirely straight-forward relation between their activity rates. QFCA provides DCE and EDCE relating the rates of directionally coupled reactions, which naturally generalize FCE given by the full coupling definition. Later on, we intend to generalize the latter example from FCE to DCE. The point that makes DCE especially suitable for this purpose is that, just like FCE, it both maintains the steady-state conditions and respects the irreversibility constraints because of the positivity of the coefficients of (2).

Suppose that in a metabolic network 𝒩= (ℳ, ℛ, *S*, ℐ), all the blocked reactions have already been removed and all the fully coupled reactions have already been merged so that there exists no more previously investigated reductions applicable to further reduce 𝒩. For any *R*_*j*_ ∈ℛ such that 𝒟_*j*_ *≠ ϕ* and (2) holds, we refer to 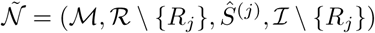 where

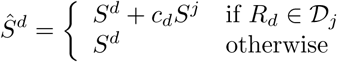

as a *DCE-induced reduction* of 𝒩 if 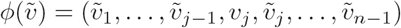, where

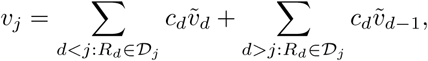

and

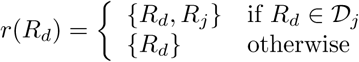

is its corresponding reduction map.

One can easily check to see that this is actually a legitimate metabolic network reduction. Therefore by applying the reduction theorem, we conclude that all the flux coupling relations of the reduced network are valid in the original network as well. Moreover, we already know which reactions are directionally coupled to *R*_*j*_. But then again, how can we tell whether *R*_*j*_ *→ R*_*i*_ or not for some *R*_*i*_ *∈*ℛ *\* {*R*_*j*_}, without undoing the changes which were applied to 𝒩 ? The following Lemma answers this question.

**Lemma 5.2.** *Let r*^-1^(*R*_*i*_) *denote the set of reactions 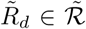 such that 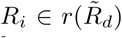. For an arbitrary pair R*_*i*_, *R*_*j*_ *∈*ℛ, *if each of the reactions in r*^-1^(*R*_*i*_) *is directionally coupled to some reaction in r*^-1^(*R*_*j*_), *then R*_*i*_ → *R*_*j*_.

*Remark.* As an immediate corollary for any arbitrary metabolic network reduction, *R*_*i*_ *→ R*_*j*_ if *r*^-1^(*R*_*i*_) *⊆ r*^-1^(*R*_*j*_).

*Proof.* If *ν*_*i*_ *≠* 0 for some fixed *ν ∈ 𝒞*, then we have

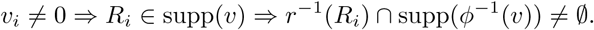

Without loss of generality, assume that 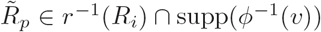. Since 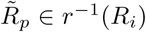, there exists 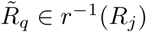 such that 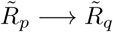. Therefore,

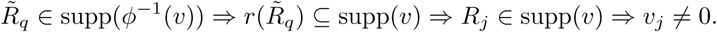

Since *ν ∈ 𝒞* was arbitrary, the proof of the lemma is complete. □

By applying the above Lemma to the DCE-induced reduction corresponding to *R*_*j*_, if all of the reactions in *r*^-1^(*R*_*j*_) are directionally coupled to *R*_*i*_, then *R*_*j*_ *→ R*_*i*_. Yet the converse is also true, because directional coupling is transitive, thus 𝒟_*j*_ *→R*_*j*_ *→ R*_*i*_. Hence in order to determine if *R*_*j*_ *→ R*_*i*_ holds in the original metabolic network, it is enough to check if *r*^-1^(*R*_*j*_) *→ R*_*i*_ holds in the DCE-reduced metabolic network. This derives all the extra flux coupling relations of 𝒩 as was promised before.

## Footnotes

1 https://mtefagh.github.io/qfca/

